# Reducing Foetal Bovine Serum Culture Conditions does not affect GPCR Signalling

**DOI:** 10.1101/2025.09.15.676239

**Authors:** Abigail Pearce, Michael Collins, Aaron Dexter, Anjana Saji, Claudia M Sisk, Emily Taylor, Yunzhu Yu, Edward Wills, Graham Ladds

**Affiliations:** Department of Pharmacology, University of Cambridge

**Author notes:** Authors contributed equally (Michael Collins, Aaron Dexter, Anjana Saji, Claudia M Sisk, Emily Taylor, Yunzhu Yu). Corresponding Author: Graham Ladds. **Declaration:** The authors have no competing interests to declare.

**Keywords:** G Protein-Coupled Receptor, Cell Signalling, Genetic Reporter, Foetal Bovine Serum

## Abstract

Animal use in research extends beyond the use of animal models to study physiology and disease. Many aspects of *in vitro* research use reagents derived from animals, most prolifically the use of foetal bovine serum (FBS) in growth media for cellular models. With the aim to reduce animal use, we investigated the effect of reduced FBS culture conditions on cell growth, as well as different stages of G protein-coupled receptor (GPCR) signalling, a wide area of research which might therefore impact many groups. We identified little differences on cell growth or GPCR signalling when reducing culture FBS percentage from 10% to 5%, using assays ranging from receptor activation to downstream transcription factors stimulation. In addition to diminishing animal use, the reduction of FBS use will also have economic and environmental benefits, which we hope will be of benefit to the wider research community.

## Introduction

Much of our understanding of modern medicine depends on research conducted using animals, with evidence of animal testing dating back to Greek scientists Aristotle and Erasistratus (1). New pharmacological treatments must be tested in animals, as *in vitro* studies cannot recapitulate the complex interactions between different organs and tissues, although researchers are employing the ‘3 Rs’, to minimise animal suffering (2). These involve replacement (e.g. use of *in vitro* models), reduction (fewer animals), and refinement (minimising pain and ensuring proper housing and care). However, these are generally only considered regarding use of live, or *ex vivo*, animals and tissue.

Animal use extends far beyond the animal house. Many reagents used routinely are sourced from animals. Most prolifically, model cells and organisms are cultured in medium that contains products of animal origin, such as the growth supplement Foetal Bovine Serum (FBS, also known as foetal calf serum, FCS), which is isolated from the bovine foetus during the slaughter of cows intended for the meat industry. Whilst it is considered a by-product, there are ethical debates regarding the ability of the foetus to feel, and there is no way to conclusively determine its suffering. As such, synthetic substitutes have been developed, although their exact composition is unknown and may also suffer from batch variability (3). There are, therefore, concerns as to their suitability, with many researchers reluctant to switch, due to the potential effect not just on cell growth, but cellular signalling and function.

One substantial area of research is the study of G protein-coupled receptors (GPCRs). GPCRs comprise the largest family of membrane proteins and are incredibly promising therapeutic targets; indeed, over 30% of all FDA approved drugs target this family (4). With growing interest, studies into the signalling of this family of proteins are increasing. GPCRs bind a range of agonists, and through coupling to intracellular G proteins, transduce these signals across cell membranes. Their signalling is complex, with 16 different G proteins genetically encoded and further potential for splice isoforms. Depending on the subfamily, these can then increase or decrease cyclic adenosine monophosphate (cAMP), mobilise Ca^2+^ from intracellular stores, and activate the Rho family of GTPases. This is not the limit of their signalling potential, with GPCR activation linked with changes in cell motility, neuronal excitability, gene expression, and even cell death. Due to its nature, the exact composition of FBS varies(5). However, it contains growth factors, lipids, and other nutrients, many of which activate GPCRs. We therefore sought to determine if a reduction in routine FBS percentage, from the recommended 10% to 5%, had any significant effect on growth of model cell lines. We then examined different signalling outputs of classical GPCRs to determine if culturing cells in lower FBS concentration had any significant effect. We determined, using transfected and endogenously expressed receptors, that the FBS percentage in growth media had no effect on GPCR signalling. Our hope is that the wider research community studying GPCRs and other receptors might use this information to give them confidence to reduce their routine FBS percentage, thereby reducing animal use.

## Results

### Reducing FBS Percentage had no effect on HEK293 cell proliferation

HEK293 cells are commonly used to measure GPCR signalling and were therefore chosen for the study, focusing on variants, HEK293A and HEK293T, used for their enhanced adherence and transfection efficiency respectively. To determine any effect on cell growth, cells were seeded at varying densities in 5% and 10% FBS, and their proliferation measured over 72 hours (Figure 1A-B). No significant differences were observed. It was also confirmed that FBS percentage had no effect on cell retention with sequential harvesting using 0.05% trypsin (Figure 1C).

**Figure 1.**
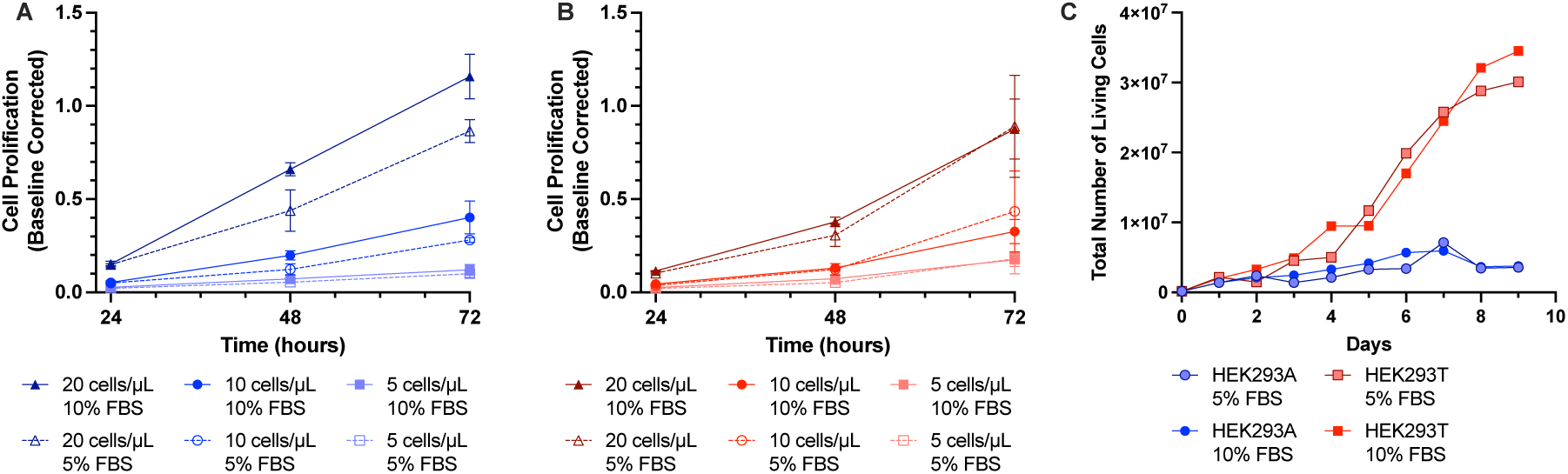
FBS Percentage has no effect on HEK293 Cell Proliferation. HEK293A and HEK293T cell proliferation was measured following culture in 5% or 10% FBS. Cells were seeded at increasing densities of 5, 10 and 20 cells/uL (A, B, C) and proliferation was assessed after 24, 48 and 72 hours. Data is mean values of n=3 performed in triplicate and corrected to absorbance measured in wells containing media alone. The total number of living cells was also measured over the course of nine days (D).

### FBS concentrations do not alter G protein-mediated signalling

We then assessed the influence of FBS concentration on G protein coupling, as measured using the bioluminescent resource energy transfer (BRET)2-based TRUPATH biosensor platform (6). BRET assays are routinely used for studying GPCR pharmacology, enabling kinetic measurements of signalling in live cells (7), and therefore interference of cell culture might influence additional outputs not studied here. We first examined any differences in the basal BRET ratio, which would be influenced by constitutive activity and transfection efficiency (Figure 2A). We used prototypical GPCRs from different G protein families; the β_2_-adrenoceptor (β_2_AR) for Gαs and the muscarinic acetylcholine receptor M3 (M_3_R) for Gα_q_. The Glucagon-Like Peptide-1 receptor (GLP-1R) was used as an example of a pleiotropic GPCR, coupling to both Gα_s_ and Gα_q_ (the latter was measured using a mutated Gα_q_ (R183Q) which displays increased activation (8)). No difference was observed in basal Gα_q_ activation, but there was an observed elevation in the basal BRET ratio for Gα_s_ for cells cultured in 10% FBS compared to 5% FBS, however it was not significant. In all cases, there was no significant effect of FBS concentration on the potency or efficacy of agonist induced G protein activation (Figure 2B-C).

**Figure 2.**
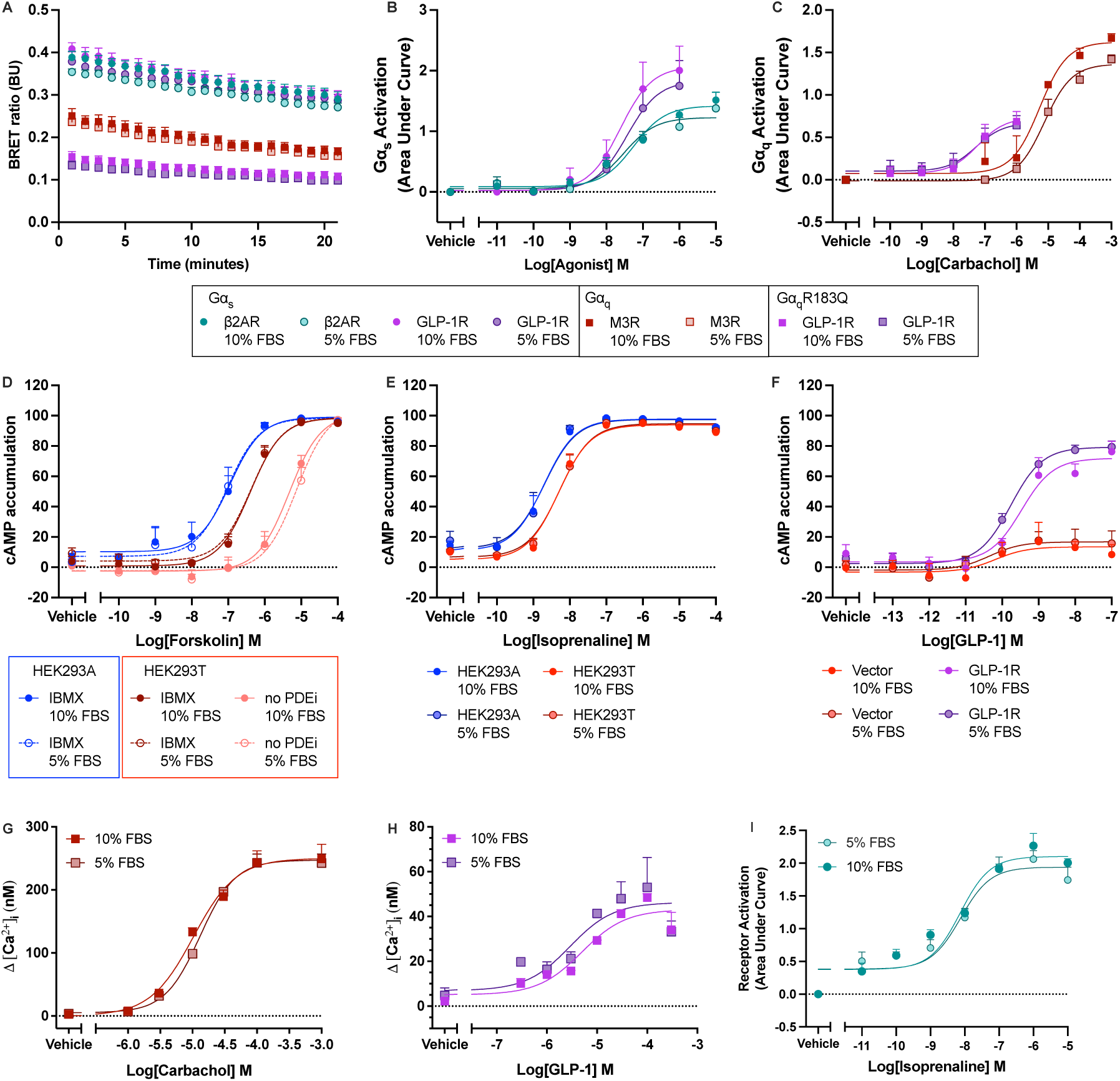
FBS growth conditions do not affect GPCR-G protein activation. (A) Basal BRET ratios for Gαs and Gαq (WT and R183Q), in HEK293T cells transfected with TRUPATH biosensors and GPCRs. (B) Gαs activation in response to stimulation of the β_2_AR with isoprenaline or the GLP-1R with GLP-1. (C) Gαq activation in response to stimulation of the M_3_R with carbachol or Gαq R183Q activation in response to GLP-1R stimulation with GLP-1. (D) Forskolin-evoked accumulation of cAMP in HEK293A or HEK293T cells, treated with in the absence or presence of IBMX. (E) Accumulation in response to isoprenaline in HEK293A and HEK293T cells incubated in 500mM IBMX. (F) HEK293T cells transfected with pcDNA3.1 (vector) or GLP-1R, stimulated with GLP-1 in the absence of PDEi. (G) Intracellular calcium measurements from HEK293A cells, stimulated with carbachol. (H) Intracellular Ca^2+^measurements from HEK293T cells transiently expressed GLP-1R, stimulated with GLP-1. (I) Activation of the β_1_AR in response to isoprenaline measured using and intramolecular BRET sensor, stably expressed in HEK293A cells. Values are shown as mean response of n repeats, where n > 2, ±SEM.

GPCRs signal via a range of second messengers following G protein activation. We therefore investigated the effects of FBS growth conditions on examples of these; cAMP accumulation and Ca^2+^ mobilisation. As these responses can often be detected at endogenously expressed receptors, we looked at the response to β_2_AR and M_3_R endogenously expressed in HEK293A and HEK293T cells, as well as the response to GLP-1R, which was transiently transfected.

We first determined that direct activation of adenylyl cyclase with forskolin was unaffected by culture conditions, using the LANCE*ultra* cAMP detection kit. Responses were determined in the absence and presence of a phosphodiesterase 4 inhibitor, rolipram, but in neither case did FBS conditions affect cAMP accumulation (Figure 2D). This was also the case for cells stimulated with agonists isoprenaline and GLP-1 (Figure E-F). Similarly, FBS culture conditions had no effect on the Ca^2+^ mobilisation of the endogenously expressed M_3_R or transiently expressed GLP-1R (Figure 2G-H).

To confirm little differences were observed at the level of the receptor, independently of any cellular signalling proteins, we examined the effects of FBS on an intramolecular biosensor of the β_1_-adrenoceptor. Receptor activation was measured through a loss of BRET between a C-terminal NanoLuc (NLuc) and mCitrine positioned within the third intracellular loop. Again, no differences were observed under different FBS culture conditions (Figure 2I).

### FBS has no effect of the sustained effects of GPCR signalling

As a downstream metric of GPCR activation, we looked at the influence of FBS culture conditions on gene expression using genetic response elements. These utilise luciferase enzymes, whose expression is controlled by a response element and its associated binding protein. We utilised two different luciferase reporters; downstream of cAMP accumulation we used cAMP Response Element (CRE) paired with NanoLuc (NLuc) (CRE-NLuc), and downstream of Ca^2+^ mobilisation we used Nuclear Factor of Activated T-Cells Response Element (NFAT) paired with (NLuc) (NFAT-Nluc). Neither basal luminescence (Figure 3A), nor the response to a cocktail of 20% FBS, 100 nM PMA, and 100 nM ionomycin were affected by the FBS condition (Figure 3B). Indeed, the responses to GPCR agonists were also unaffected, consistent with the effects on second messenger signalling (Figure 3C-E).

**Figure 3.**
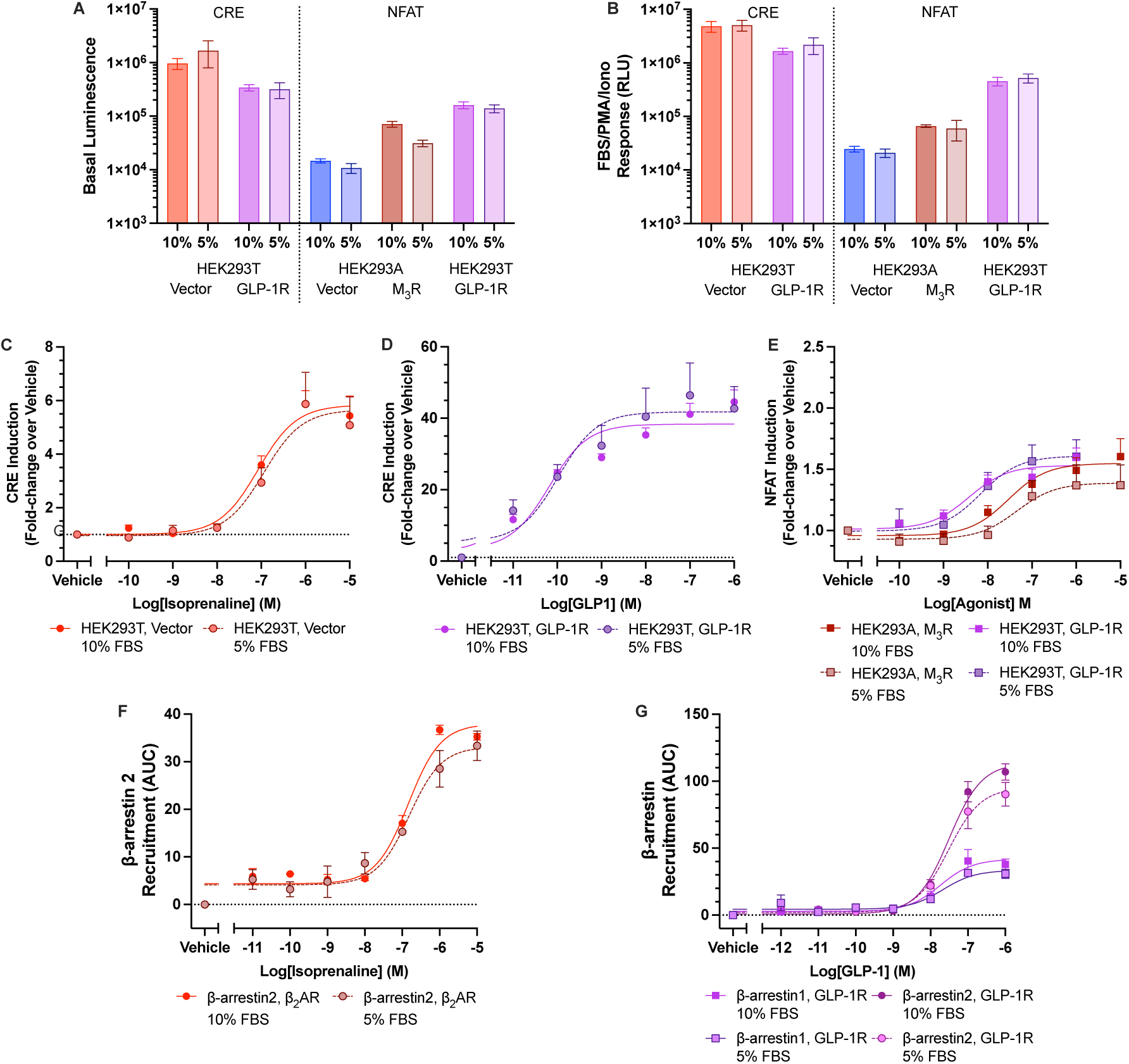
FBS Growth conditions do not alter sustained signalling or desensitisation. (A) Basal luminescence measured in HEK293T and HEK293A cells transfected with either CRE-NLuc or NFAT-NLuc and and either GLP1-R, M_3_R, or vector. RLU values for the mixture of 20% FBS, 100 nM phorbol-12-myristate-13-acetate (PMA), 100 nM ionomycin across the same cell groups expressing CRE-NLuc or NFAT-NLuc (B). Fold-change induction of CRE-NLuc responses were measured following stimulation of HEK293T cells with isoprenaline (C) or GLP-1, in cells expressing GLP-1R (D). (E) NFAT-NLuc responses were measured in response to stimulation with carbachol, in HEK293A cells transfected with the M_3_R, and in HEK293T cells expressing GLP-1R, stimulated with GLP-1. (F) β-arrestin-2 recruitment to β_2_AR in response to isoprenaline. (G) β-arrestin1 and 2 recruitment to GLP-1R in response to GLP-1. Responses were measured in HEK293T cells transfected with RIT-Venus, β-arrestin-Nluc, and the appropriate GPCR. Values are plotted as mean ± SEM of n experiments performed in duplicate, where n>3.

Lastly, we examined the effects on receptor desensitisation by looking at β-arrestin recruitment. These are recruited to the active receptor, blocking G protein coupling. As prototypical members of Class A and Class B1 GPCRs, we focused on the β_2_AR and the GLP-1R. To avoid using tagged receptors, recruitment of a NLuc-tagged arrestin to the plasma membrane was measured through an increase in BRET with Venus-tagged RIT, a plasma membrane bound GTPase. In line with the results on G protein coupling, β-arrestin recruitment was also unaffected by FBS culture conditions (Figure 3F-G).

## Discussion

With established protocols, it is hard to change conditions, and alterations are often kept within a group as they are not deemed significant to the wider community. We sought to determine if these conditions could be altered, without a loss of cell viability or changes to signalling of our target of interest, GPCRs. These changes were motivated by a desire to reduce the usage of animal products, but also present economic and environmental benefits with the increasing price of FBS and concerns around sustainability respectively. We found no statistical differences in the signalling of GPCRs, looking across G protein activation to regulation of gene transcription. Whilst these experiments were carried out across a finite set of receptors and signalling outputs, they give us confidence that, moving forward, growth media FBS percentage can be reduced. However, it is recommended to conduct small screening experiments when studying your own receptors, as they might be more directly influenced by the different components within FBS, such as receptors activated by fatty acids or growth factors.

## Methods

### Cell lines

HEK293A cells (a gift from David Yule) and HEK293T cells were cultured in Dulbecco’s modified eagle medium (DMEM)/F12 Glutamax, supplemented with 1% antibiotic-antimycotic solution, and either 5% or 10% foetal bovine serum (FBS). Cells were thawed in the appropriate percentage and cultured for at least 7 days before the start of any experiments.

### Quantification of Cell Proliferation

HEK293A and HEK293T cells were seeded into 96-well plates (Greiner), at 5, 10, or 20, cells/μL (100μL total), and cultured in media containing 5% or 10% FBS. Media alone was also added to wells as a negative control. Plates were incubated for 24, 48 and 72 hours. 5uL of Cell Counting Kit-8 (CCK-8, Sigma Aldrich) was added per well and incubated for three hours. Absorbance was then measured at 460nm using the SpectraMax iD3 microplate reader (Molecular Devices).

HEK293A and HEK293T cells grown in media containing 5% or 10% FBS were also seeded at a low density (50,000/ml) into a 75cm^2^ tissue culture-treated flask (corning). Cells were passaged using 0.05% trypsin (Sigma Aldrich) daily, and counted using the Countess (Thermo-Fisher), using trypan blue to distinguish live cells.

### G Protein Dissociation Assay

G Protein dissociation was measured using the TRUPATH biosensor platform. HEK293T cells were transfected with receptor and Gα-Rluc8, Gγ-GFP2, and Gβ at a 1:1:1:1 ratio, using polyethyleneimine, at a 1:6 ratio (*w/v*) in 150mM NaCl. After 24 hours, cells were seeded into poly-L-lysine (PLL)-coated white 96-well plates and cultured for a further 24 hours. On the day of assay, cells were washed with Hank’s Balanced Salt Solution (HBSS) before addition of HBSS supplemented with 20 mM HEPES, 0.1% Bovine Serum Albumin (BSA), and 5μM (Nanolight Technology, USA). Agonist (1 mM to 0.01 nM) was then added and plates recorded for 20 minutes on a PHERAstar (BMG Labtech, UK) plate reader at 60 second intervals, using a BRET2 optical module. The BRET ratio was calculated as λ_515nm_/λ_400nm_.

### Mobilisation of intracellular Ca^2+^

Carbachol-evoked changes in intracellular calcium were recorded from cell populations of HEK293A cells grown in black-walled PLL-coated 96-well plates. HEK293T cells grown to 80% confluency in a 6-well plate, were transfected with GLP-1R using Transit-LT1 according to manufacturer instructions (Mirus, USA). Following 24 hours, cells were seeded into black-walled PLL-coated 96-well plates.

Cells were incubated with 2µM Fluo-8 (AAT-Bioquest), (1h at 20 °C) in HEPES-buffered saline (HBS - NaCl 135mM, KCl 5.9mM, MgCl_2_ 1.2mM, CaCl_2_ 1.5mM, glucose 11.5mM, and HEPES 11.6mM, pH 7.3) with 0.02% w/v pluronic acid. Fluorescence was recorded using a FlexStation 3 fluorescence plate-reader (Molecular Devices) and calibrated to intracellular calcium as described (9).

### Quantification of cAMP accumulation

cAMP accumulation was measured using the LANCE*ultra* cAMP detection kit (Revvity) as previously described. Briefly, HEK293A and HEK293T cells were harvested, and resuspended in phosphate buffered saline (PBS), supplemented with 0.1% BSA, at 200 cells/μL in the presence of IBMX, or 100 cells/μL in the absence of IBMX. Cells were plated on a 384-well OptiPlate in 5μL aliquots and stimulated with agonist for 30 minutes at room temperature, before detection reagents were applied. Plates were read using a Mithras LB 940 multimode microplate reader (Berthold Technologies, Germany) using the TRF module. Responses were normalised to the minimum and maximum detection, determined using the cAMP standard, to account for differences in constitutive activity.

### Intramolecular GPCR Activation Biosensor

An intramolecular biosensor for the thermostabilised wild turkey β1 adrenoceptor (10), with an mCitrine inserted into intracellular loop 3, and a NLuc at the C terminus, after helix 8, was provided by Dr Suleiman Al-Sabah (Kuwait University). HEK293A cells were transfected with the biosensor and cultured with 1000μg/mL of G418 in DMEM/F12 supplemented with 10% FBS to generate a stable cell line. Frozen cell aliquots were thawed and subsequently cultured in 5 or 10% FBS prior to experiments. Cells were harvested and seeded in white 96-well plates at 25,000 cells per well 24 hours before assay. On the day of assay, media was removed and cells incubated in PBS supplemented with 0.1% BSA, 0.9mM CaCl_2_ and 0.4mM MgCl_2_, and NanoGlo® (1 in 10000 dilution) for five minutes. Cells were stimulated and the plate read immediately using the CLARIOstar Plus® microplate reader (BMG Labtech, UK), using filters at 410nm and 530nm for 30 minutes. The BRET ratio was calculated as λ_530_/λ_410_ and the area under curve for the response period used to generate dose response curves.

### Report of Downstream Signalling via Luminescent Gene Expression

The signalling pathways downstream of cAMP and Ca^2+^ mobilization were measured via CRE and NFAT response elements paired with luminescent reporter genes. HEK293A and HEK293T cells were transfected with NanoLuc Luciferase (NLuc) genetic reporters, whose expression is dependent on CRE (CRE-Nluc) or NFAT (3xNFAT-Nluc), and M3R or GLP1-R (or pcDNA3.1(+) in cases of testing endogenous receptor) in a 1:1 ratio. Luminescent reporter plasmids consisting of a pause site, minimal promoter, Nano-luciferase, PEST sequence and bgH Poly(A) region were synthesised (Twist Bioscience). Using NheI and HindIII, CRE and NFAT-RE transcriptional response elements were subcloned from commercially available pNL[CRE-NLuc-PEST-Hygro] and pNL[NFAT-RE-NLuc-PEST-Hygro] constructs (Promega, UK), respectively. After remaining in transfection media for 24 hours, cells were seeded in PLL-coated 96-well plates at a concentration of 000 cells/uL and volume of 100 uL in DMEM/F12 (1:1) with GlutaMAX™ without phenol red (ThermoFisher Scientific, UK). At time of plating, CRE reporter cells underwent serum starvation and both 5% and 10% FBS CRE cells were reduced to 0.5% (v/v) FBS. NFAT reporter cells remained at either 5% or 10% FBS in the 96-well plate. Cells were stimulated in media containing 0.1% BSA and maintained at 37°C and 5% CO_2_ for 3 hours for NFAT and 6 hours for CRE. After this incubation period, media was replaced with 50uL of 1x lysis mixture (Passive Lysis Buffer (PLB), Promega, UK) containing substrate. A 1:1000 dilution of NanoGlo® (furimazine; Promega, UK) was used for NFAT, and a 1:500 dilution of coelenterazine-H (Nanolight Technology, USA) was used for CRE. Plates were shaken in this lysis mixture for 10 minutes before being measured for endpoint luciferase activity via unfiltered luminescence measurement with a CLARIOstar Plus® (BMG Labtech, UK) with Enhanced Dynamic Range. Dose response curves were generated by calculating the fold change in luminescence (luminescence of ligand divided by luminescence of vehicle).

### Bystander assay for β-arrestin recruitment

HEK293T cells were transfected with 1.2μg of total DNA containing the receptor of interest, N terminally NLuc-tagged β-arrestin1 or 2 (Nluc-arrestin) and RIT-Venus (donated by Luke Pattinson, University of Cambridge) at a 1:1:1 ratio. 24 hours post-transfection, cells were reseeded onto white 96 well plates at a density of 5×10^4^ cells/well in 100μl growth medium and left to incubate overnight. After 24 hours, media was removed, and cells were washed with PBS. Serial dilutions of ligands were added to assay buffer (PBS supplemented with 0.1% BSA, 0.9mM CaCl_2_ and 0.4mM MgCl_2_) from 0.01nM to 10μM. Cells were incubated in buffer supplemented with 0.02% NanoGlo for 5 minutes. Cells were stimulated and the plate read immediately using the CLARIOstar Plus® microplate reader (BMG Labtech, UK), using filters at 410nm and 530nm for 30 minutes. The BRET ratio was calculated as λ_530_/λ_410_ and the area under curve for the response period used to generate dose response curves.

### Data Analysis

Data was analysed using GraphPad Prism 10. Dose response curves were calculated using the three-parameter log[agonist] vs response model, to give pEC_50_ and E_max_ values. Statistical significance was tested using Two-Way ANOVA or Student’s T test where appropriate, with p<0.05 considered statistically significant.

## Abbreviations

FBS: Foetal Bovine Serum
FCS: Foetal Calf Serum
GPCR: G Protein-Coupled Receptor
cAMP: cyclic adenosine monophosphate
HEK293: human embryonic kidney 293
β_1_AR: β1-adrenoceptor
β_2_AR: β2-adrenoceptor GLP-1, glucagon-like peptide-1
GLP-1R: GLP-1 receptor
M_3_R: muscarinic acetylcholine receptor M3
NLuc: NanoLuc
CRE: cAMP response element
NFAT: nuclear factor of activated T cells;

## Author Contributions

A.P., M.C., A.D., A.S., C.M.S., E.T., Y.Y., performed experiments and completed data analysis. A.P. and E.W. synthesised constructs, and developed methodologies. A.P. conceived the experiments and wrote the manuscript. G.L. provided supervision and funding. All authors proof-read and provided feedback on the manuscript.

## Acknowledgements

Work was funded by the Biotechnology and Biological Sciences Research Council (BBSRC; BB/Y513817/1 to G.L.; BB/W014831/1 to A.P.). We thank the Harding Distinguished Postgraduate Scholars Programme and the David James Trust fund for funding A.D. M.C is funded by a BBSRC iCase studentship (BB/Z516454/1) in partnership with Syngenta. E.T. is funded by the Leverhulme Trust (RPG-2023-130). C.M.S was funded by a Cambridge Trust International Scholarship in conjunction with the Gonville and Caius Stanley Elmore PhD Studentship. E.W. is funded by an Engineering and Physical Sciences Research Council iCase studentship co-funded with AstraZeneca (EP/X015785/1, E.W.). G.L. is a Royal Society Industry Fellow (INF/R2/212001). We thank Dr Suleiman Al-Sabah for providing the intramolecular biosensor.

## References

1. M. Sántha, Biologia futura: animal testing in drug development—the past, the present and the future. Biol Futur 71, 443–452 (2020).

2. W. T. Poh, J. Stanslas, The new paradigm in animal testing – “3Rs alternatives.” Regulatory Toxicology and Pharmacology 153, 105705 (2024).

3. J. van der Valk, Fetal bovine serum—a cell culture dilemma. Science (1979) 375, 143–144 (2022).

4. A. S. Hauser, M. M. Attwood, M. Rask-Andersen, H. B. Schiöth, D. E. Gloriam, Trends in GPCR drug discovery: new agents, targets and indications. Nat Rev Drug Discov 16, 829–842 (2017).

5. H. Barosova, et al., Inter-laboratory variability of A549 epithelial cells grown under submerged and air-liquid interface conditions. Toxicology in Vitro 75, 105178 (2021).

6. R. H. J. Olsen, et al., TRUPATH, an open-source biosensor platform for interrogating the GPCR transducerome. Nat Chem Biol 16, 841–849 (2020).

7. A. Pearce, et al., Quantitative approaches for studying G protein-coupled receptor signalling and pharmacology. J Cell Sci 138 (2025).

8. D. Safitri, et al., Cancer-Associated Mutations Enhance The Sensitivity Of The Trupath GαQ/11 System. bioRxiv 2022.09.01.506210 (2022). 10.1101/2022.09.01.506210.

9. E. Pantazaka, E. J. A. Taylor, W. G. Bernard, C. W. Taylor, Ca2+signals evoked by histamine H1 receptors are attenuated by activation of prostaglandin EP2 and EP4 receptors in human aortic smooth muscle cells. Br J Pharmacol 169, 1624–1634 (2013).

10. A. J. Y. Jones, et al., Binding kinetics drive G protein subtype selectivity at the β1-adrenergic receptor. Nat Commun 15, 1334 (2024).

